# Shotgun proteomics of SARS-CoV-2 infected cells and its application to the optimisation of whole viral particle antigen production for vaccines

**DOI:** 10.1101/2020.04.17.046193

**Authors:** Lucia Grenga, Fabrice Gallais, Olivier Pible, Jean-Charles Gaillard, Duarte Gouveia, Hélène Batina, Niza Bazaline, Sylvie Ruat, Karen Culotta, Guylaine Miotello, Stéphanie Debroas, Marie-Anne Roncato, Gérard Steinmetz, Charlotte Foissard, Anne Desplan, Béatrice Alpha-Bazin, Christine Almunia, Fabienne Gas, Laurent Bellanger, Jean Armengaud

**Author notes:** These two co-authors should be considered as co-first authors (Lucia Grenga, Fabrice Gallais). Corresponding author: Jean Armengaud, CEA-Marcoule, DRF-Li2D, Laboratory “Innovative technologies for Detection and Diagnostics”, BP 17171, F-30200 Bagnols-sur-Cèze, France.

## Abstract

Severe acute respiratory syndrome-related coronavirus 2 (SARS-CoV-2) has resulted in a pandemic and continues to spread quickly around the globe. Currently, no effective vaccine is available to prevent COVID-19 and an intense global development activity is in progress. In this context, the different technology platforms face several challenges resulting from the involvement of a new virus still not fully characterised. Finding of the right conditions for virus amplification for the development of vaccines based on inactivated or attenuated whole viral particles is among them. Here, we describe the establishment of a workflow based on shotgun tandem mass spectrometry data to guide the optimisation of the conditions for viral amplification. In parallel, we analysed the dynamic of the host cell proteome following SARS-CoV-2 infection providing a global overview of biological processes modulated by the virus and that could be further explored to identify drug targets to address the pandemic.

## Introduction

SARS-CoV-2 belongs to the B lineage of the beta‐coronaviruses and is closely related to the SARS‐CoV virus [1,2]. It is the causative agent of COVID-19, a severe acute respiratory syndrome that spread world-wide within a few weeks starting on December 2019 in Wuhan [3]. The major four structural genes encode the nucleocapsid protein (N), the spike protein (S), a small membrane protein (SM) and the membrane glycoprotein (M) with an additional membrane glycoprotein (HE) occurring in the HCoV‐OC43 and HKU1 beta‐coronaviruses [3].

Based on the speed at which the outbreak of COVID-19 has developed, SARS-CoV-2 appears to spread easily in the human population. The reproductive number (*R*_0_) of the virus is currently thought to be around 3, suggesting the potential for sustained human-to-human transmission that appears to be through respiratory droplets and potentially a fecal-oral route [4]. In this pandemic situation, one of the outstanding questions concerns the possibility to contain the spread of SARS-CoV-2 and its persistence in the human population. Social distancing policy, lock-down and other containment measures have been worldwide implemented to slow down the spread. Current roadmaps to lifting the restrictions rely on the deployment of effective diagnostics, therapies, and eventually on the development of an effective vaccine.

Several platforms are being used to develop vaccines against SARS-CoV-2, including spike subunit, DNA, RNA, whole-virion, and nanoparticle vaccines. Most successful antiviral vaccines employ inactivated or attenuated whole viral particles as vaccine antigen and depend on the induction of neutralizing antibodies [5,6] against structural proteins of the virus. However, virus yields from the dedicated cell culture systems could be relatively low compared to quantities envisioned to be required for massive vaccine production. In addition, the production campaigns are time-consuming and highly demanding due to the danger of working with these pathogens, and thus optimization of the production of whole viral particle antigen is of utmost interest for vaccines. Concomitantly with vaccine development, a better understanding of how the host responds to SARS-CoV-2 infection may help direct further therapeutic avenues.

Multiple proteomics strategies have been shown insightful for better understanding of coronavirus structure and its molecular mechanisms of infection. Tracheal tissues of chicken infected with infectious bronchitis coronavirus were analyzed by 2D-DIGE and MALDI-TOF tandem mass spectrometry to establish the host response [7]. Vero cells infected with porcine epidemic diarrhea virus (PEDV) were analyzed by shotgun proteomics [8]. Different PEDV coronavirus strains were compared with an iTRAQ-labeling quantitative approach showing differences of inflammatory cascade eliciting [9]. The dynamics of the host proteins triggered by specific overexpressed coronavirus genes was also established [10]. While no literature is yet available on the proteomics characterization of SARS-CoV-2 virus, several studies of interest have been recently submitted and should be soon available [11,12,13].

Here, we describe the establishment of a workflow based on shotgun tandem mass spectrometry data that in addition to gaining more basic information about SARS-CoV-2 infection aims at guiding the optimisation of the conditions for whole viral particle antigen production and aiding SARS-CoV-2 vaccine development.

## Materials and Methods

### Cell culture and Virus preparation

Vero E6 (ATCC, CLR-1586) cells were cultured at 37°C in 9% CO_2_ in Dulbecco’s modified Eagle’s medium (DMEM, Gibco™, ThemoFisher) supplemented with 5% fetal calf serum (FCS) and 0.5% penicillin–streptomycin. Cells were passaged by trypsinization every 2 days. The SARS-CoV-2 strains 2019-nCoV/Italy-INMI1 (Genbank MT066156) was provided by the Lazzaro Spallanzani National Institute of Infectious Diseases (Rome, Italy) via the EVAg network (European Virus Archive goes global). SARS-CoV-2 stocks used in the experiments had undergone two passages on Vero E6 cells and were stored at −80°C. Virus titer was determined by standard plaque assay (1×10^7^ pfu/ml).

### Infection

For the kinetic, 1×10^6^ Vero cells seeded into 25 cm^2^ flasks were grown to cell confluence in 5 mL DMEM supplemented with 5% FCS and 0.5% penicillin–streptomycin for one night at 37°C under 9% CO_2_. They were infected at two multiplicities of infection (MOI): 0.01 and 0.001. Cells were harvested at 1, 2, 3, 4, and 7 days post infection (dpi). Supernatants of SARS-CoV-2 infected cells were saved for plaque assay titration to confirm production of infectious viral particles. Infected Vero cells were microscopically observed for cytopathic effect (CPE) at the same time points.

### Quantification of viral RNA by qRT-PCR

SARS-CoV-2 RNA from cell culture supernatant samples and supernatant plus cells was isolated using the NucleoSpin RNA Virus, Mini kit for viral RNA from cell-free fluids (Macherey Nagel) according to the manufacturer’s instructions. RNA was subjected to qRT-PCR analysis using the SuperScript III Platinum One-Step qRT-PCR Kit (ThermoFisher) and a CFX96 Touch Real-Time PCR Detection System Thermal Cycler (BioRad). Primers targeting IP2 and IP4 (RdRp) were as recommended [14]: nCoV_IP2-12669Fw (ATGAGCTTAGTCCTGTTG) nCoV_IP2-12759Rv (CTCCCTTTGTTGTGTTGT) nCoV_IP2-12696bProbe(+) (AGATGTCTTGTGCTGCCGGTA [5’]Hex [3’]BHQ-1) and nCoV_IP4-14059Fw (GGTAACTGGTATGATTTCG) nCoV_IP4-14146Rv (CTGGTCAAGGTTAATATAGG) nCoV_IP4-14084Probe(+) (TCATACAAACCACGCCAGG [5’]Fam [3’]BHQ-1), respectively, using 0.4 µM per reaction. Standard curves were created using in vitro transcribed RNA derived from strain BetaCoV_Wuhan_WIV04_2019 (EPI_ISL_402124). The transcript contains the amplification regions of the RdRp and E gene as positive strand. Each microtube contains 1011 copies of target sequences diluted in presence of yeast tRNAs, and lyophilised. Mean and standard deviation were calculated for each group (n=3).

### Sample preparation for mass spectrometry

At each of various time points, SARS-CoV-2 -infected Vero cells were washed twice with 5 ml of PBS to remove media and FCS used to culture the cells. A volume of 1.5 ml of PBS was added to the washed cells before harvesting. The virus was inactivated and the cells lysed by autoclaving the samples at 125°C for 40 min. Proteins were precipitated by adding cold trichloroacetic acid to a final concentration of 10% (w/v). After an incubation of 5 min at 4 °C, the precipitated material was recovered by centrifugation at 16000 xg for 10 min. The proteins in the resulting pellets were then dissolved in 100 μL LDS 1X (Lithium dodecyl sulfate) sample buffer (Invitrogen) and supplemented with 5% beta-mercaptoethanol (vol/vol) before sonication with a Hielscher UP50H disruptor operated for 20 sec at 60% amplitude with 0.25 sec impulsions and then incubation for 5 min at 99°C. A 20 µl aliquot of a 1/8 dilution in LDS 1X (invitrogen) of each sample was loaded on NuPAGE 4–12% Bis-Tris gel and subjected to short SDS-PAGE migration. The proteins were stained for 5 min with Coomassie SimplyBlue SafeStain (Thermo Fisher Scientific) prior in-gel trypsin proteolysis performed as described in Hartmann et al. [15].

### Liquid chromatography-mass spectrometry

Peptides were identified using an ultimate 3000 nano-LC system (Thermo Fisher Scientific) coupled with a Q-Exactive HF mass spectrometer (Thermo Fisher Scientific). Peptides were desalted on a reversed-phase PepMap 100 C18 μ-precolumn (5 μm, 100 Å, 300 μm i.d. × 5 mm, Thermo Fisher Scientific) before peptide separation on a nanoscale PepMap 100 C18 nanoLC column (3 μm, 100 Å, 75 μm i.d. × 50 cm, Thermo Fisher Scientific) at a flow rate of 0.2 μL per min using a 120 min gradient comprising 100 min from 4% to 25% of solvent B and 20 min from 25% to 40% of solvent B (solvent A consisted in 0.1% formic acid in water, solvent B was 80% acetonitrile, 0.1% formic acid in water). The mass spectrometer was operated in Top20 mode. Full MS were acquired from 350 to 1,500 m/z and the 20 most abundant precursor ions were selected for fragmentation with 10 s dynamic exclusion time. Ions with 2+ and 3+ charge were selected for MS/MS analysis. Secondary ions were isolated with a window of 2.0 *m/z*.

### MS/MS Data Interpretation and label free protein quantification

The MS/MS spectra recorded on each sample were assigned to peptide sequences using the Mascot Server 2.5.1 (Matrix Science). A database (20,585 sequences; 10,847,418 residues) containing the UniProt *Chlorocebus* sequences (downloaded March 2020) and the Italy-INMI1 SARS-CoV-2 protein sequences was queried after a first analysis to remove contaminant spectra against an in-house ‘common contaminants’ database (384 sequences; 187,250 residues) encompassing 361 contaminants classically observed in proteomics (cRAP + additional contaminants) and 23 *Bos taurus* sequences corresponding to the most abundant proteins from foetal calf serum present in the cell culture medium [16]. Peptide-to-MS/MS spectrum assignation was done with the following parameters: full trypsin specificity, maximum of two missed cleavages, mass tolerances on the parent ion of 5 ppm and 0.02 Da on the MS/MS, static modification of carbamidomethylated cysteine (+57.0215), and oxidized methionine (+15.9949) and deamidation of asparagine and glutamine (+0.984016) as dynamic modifications. Mascot DAT files were parsed using the Python version of Matrix Science msparser version 2.5.1 with function ms_peptidesummary. Peptide-to-Spectrum Matches (PSMs) with the expectation values corresponding to 1% False Discovery Rate (FDR) were validated using the MASCOT homology threshold option. Multiple PSMs per MS/MS spectra were allowed in case of ion scores higher than 98% of the top ion score. Proteins were grouped if they shared at least one peptide, and in each group label-free quantification was based on PSM counts for each protein following the principle of parsimony. Proteins identified by one or more specific peptides were retained for the analysis (protein FDR 1%).

### MS/MS data repository

The mass spectrometry proteomics data have been deposited to the ProteomeXchange Consortium via the PRIDE [17] partner repository with the dataset identifier PXD018594 and 10.6019/PXD018594.

### Data analysis

Principal component analysis was done as previously described [18]. Co-expression cluster analysis was obtained using the Bioconductor R package coseq v1.5.2 [19]. The protein abundance matrix was used as an input in coseqR. Log CLR-transformation was applied to the matrix to normalize the abundance of proteins and the K-means algorithm was chosen to detect the co-expressed clusters across the different time points. The K-mean algorithm was repeated 20 times in order to determine the optimal number of clusters. The resulting number of clusters in each run was recorded, and the most parsimonious cluster partition was selected using the slope heuristics approach. Both the PCA and the coseq analysis were performed after removal of proteins with spectral counts lower than three (1402 host protein groups retained). Finally, proteins assigned to the different clusters were retained for cluster visualization and gene ontology (GO)-enrichment analysis per each cluster. Statistically enriched (FDR ≤ 0.05) GO terms on proteins that are differentially expressed between pairwise samples or on proteins assigned to each co-expression cluster were identified using Metascape [20]. The most statistically enriched GO terms were visualized in ggplot2 [21].

## Results

### Profiling of virus production by tandem mass spectrometry

To understand the dynamics associated with SARS-CoV-2 infection and determine optimal conditions for whole-viral particle antigen production we infected Vero E6 cells with SARS-CoV-2 at two multiplicity of infection (MOI 0.01 and 0.001) and monitored the kinetics of the infection by means of tandem mass spectrometry over several days (Figure 1). Overall, we identified 3220 Vero cell proteins and 6 SARS-CoV-2 proteins with 27388 and 94 unique peptides (FDR below 1%), respectively (Table S1). Among the identified viral proteins, three (N, S, M) out of the four encoded by the viral genome were structural proteins while three were non-structural ones. In particular, out of the sixteen non-structural proteins we identified peptides corresponding to the ORF1a papain-like protease (PLpro) / 3C-like protease (3CLpro) and to the accessory proteins encoded by ORF3a and ORF7a. These protein sequences were covered with 40 (N), 29 (S), 7 (M), 13 (ORF1ab), 4 (ORF3a), and 1 (ORF7a) distinct peptides. The sequence coverage of these polypeptides depends on their abundance, size, and lysine and arginine residues occurrence, the ORF7a protein with 121 residues being logically poorly detected compared to the others. Based on the sets of peptides shared by the different proteins, 2984 protein groups were identified, for which a total of 457,111 PSMs were assigned. The viral proteins represented 1.4% of these PSMs.

**Figure 1.**
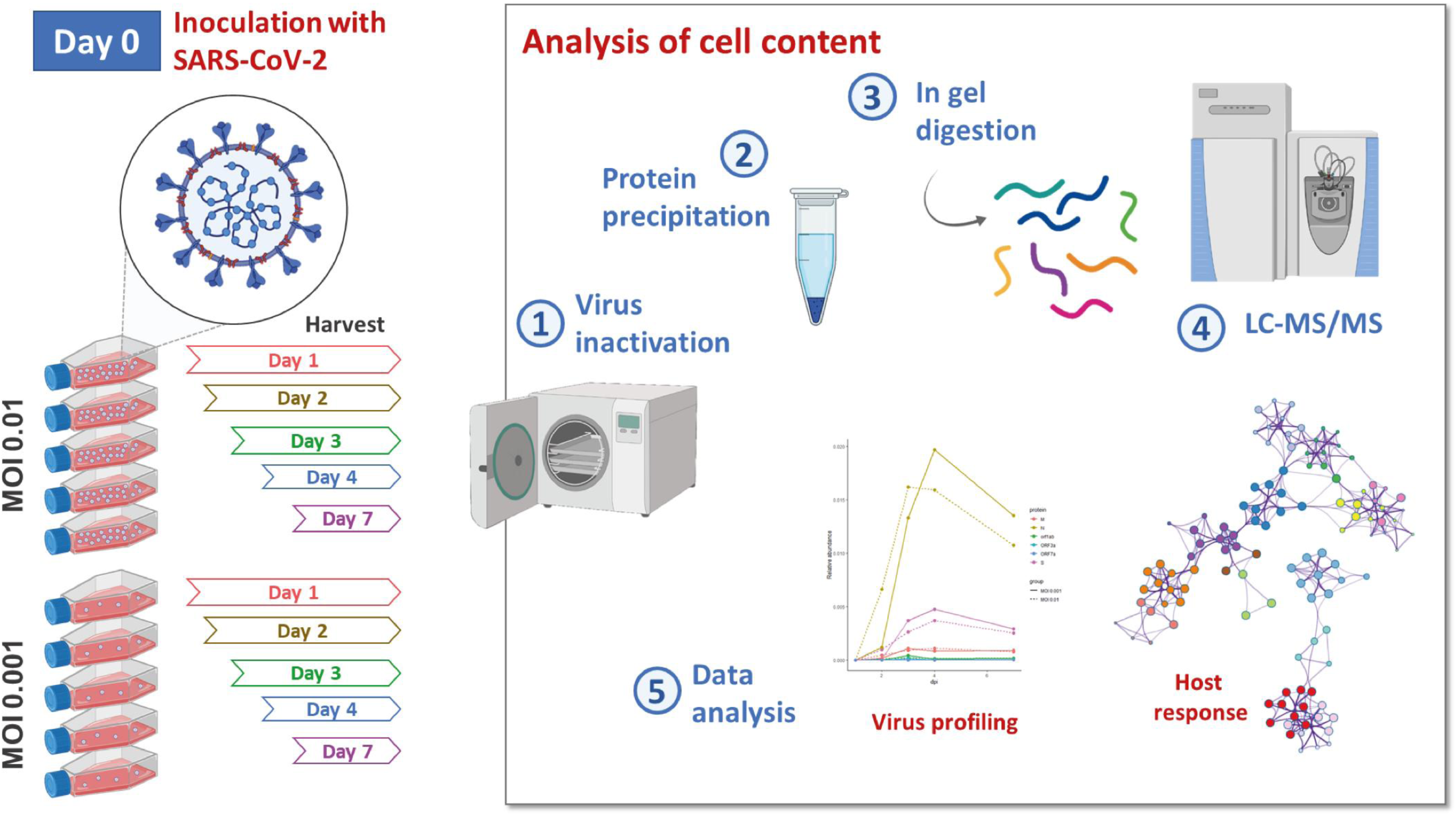
Schematic representation of the experimental design. Vero E6 cells were infected at Day 0 with SARS-CoV-2 at two multiplicity of infection (MOI 0.01 and 0.001). The kinetics of the infection was monitored by means of tandem mass spectrometry over several days. The main steps and the output of the analysis are highlighted.

The dynamics of viral protein levels across time points indicated that SARS-CoV-2 protein synthesis increased continuously after infection with a peak registered around day 3 post infection. Viral amplification was slightly delayed in cells infected at a MOI of 0.001 (Figure 2A). SARS-CoV-2 N, S, and M were consistently among the most abundant proteins detected. Their respective ratios were relatively constant along time. Decline in abundance of viral proteins registered at 7 days post infection was in accordance to notable cytopathic effect following intense virus replication observed here for each culture at this time-point, as previously reported [22].

**Figure 2.**
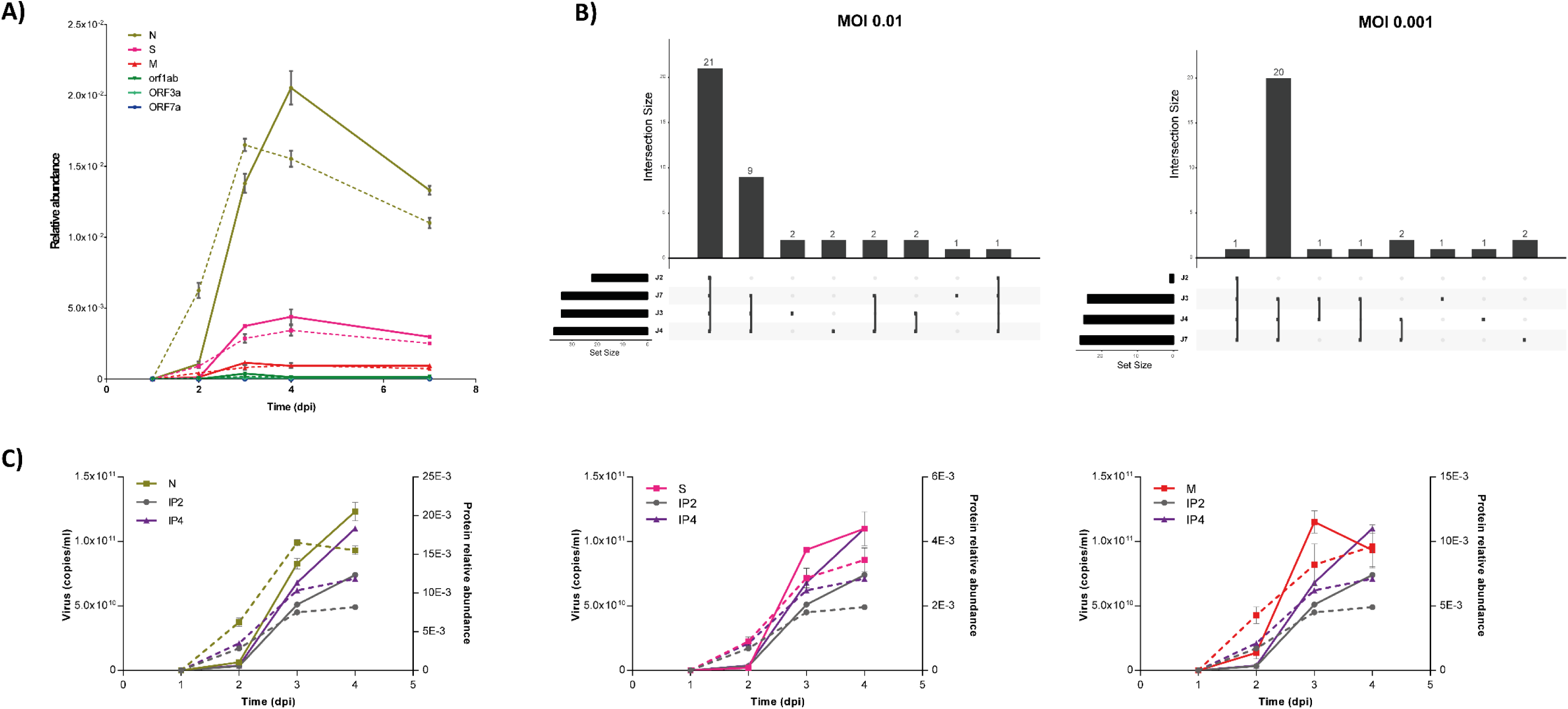
Virus profiling by tandem mass spectrometry. **(A)** Kinetic of viral production by LC-MS/MS. Relative abundance at each time points represents the mean ± standard deviation of the technical replicates at the two MOI tested. Dashed curves indicate results obtained at MOI 0.01 while solid curves refer to MOI 0.001. **(B)** Comparison between viral peptides identified at the different time points. Sets of intersections are visualized using the UpSet matrix layout and plotted horizontally. Each column corresponds to an exclusive intersection that contains the elements of the sets represented by the dark circles. Sets are represented by the different peptides assigned to viral proteins at each time point. J2, J3, J4 and J7 refer to the different time points analysed: Day 2, 3, 4 and 7, respectively. **(C)** Correlation between viral counts (copies/mL) obtained by qRT-PCR and the abundance of viral proteins measured by LC-MS/MS across time points. Dashed curves indicate results obtained at MOI 0.01 while solid curves refer to MOI 0.001.

As expected, a larger variety of peptides was found for overrepresented proteins. Figure 2B shows the distribution of this diversity across the time points for the three most abundant viral proteins. Interestingly, increase of protein levels relied on increasing abundance of the same set of peptides with only few new sequences registered at the peak of viral production compared with an early time point.

To evaluate to what extent virus profiles obtained by tandem mass spectrometry reflected virus production we measured SARS-CoV-2 RNA molecules by quantitative PCR analysis across the same time points (Figure 2C). Variations in most abundant viral protein yields reflected variation in the number of SARS-CoV–2 RNA molecules confirming that LC-MS/MS with label-free quantitation can be applied to monitor SARS-CoV-2 infection kinetics.

### Characterization of host cell protein dynamics upon SARS-CoV-2 infection

Next, we characterized changes in cellular protein networks upon SARS-CoV-2 infection at the level of total protein abundance. Dimension reduction by principal component analysis (PCA) showed that early and late time points were distinctively distributed along principal components 1 and 2, with replicates clustering closely. This was observed for both MOI. Interestingly, a degree of overlap was observed for time points J3 and J4 corresponding to the peak of viral protein abundance implying a smaller evolution of the host proteome between these time points (Figure 3A). Addition of the viral proteins measured at each time point did not affect samples separation. To elucidate the host response during virus amplification, we performed a co-expression cluster analysis to determine host proteins showing similar profiles over time (Figure 3B). Functional enrichment analysis performed on the members of each cluster provided a global overview of biological processes (Figure 4A-C). At MOI 0.01, clusters 2 and 5, with 345 and 198 detected protein members, respectively, showed a close similarity of the statistically enriched terms, with functions and pathways related to the viral life cycle among the most represented ones. Besides, as revealed by functional interaction network analysis even cluster-specific enriched functions like membrane trafficking and protein pre-processing in endoplasmatic reticulum and regulation of mRNA processing/splicing via spliceosome, respectively, were highly interconnected (Figure 4A-B). Also, an overlap of enriched biological functions was observed for clusters 3 and 6, with 16 and 10 representatives, respectively, characterized by a specific increase in the abundance of clustered proteins 3 days post-infection, with proteins involved in cornification and ECM regulators (Extra Cellular Matrix) among the top 20 significantly enriched terms (Figure 4A). Four distinct expression profiles were identified at MOI 0.001 (Figure 3B). Comparative analysis and inference of enriched biological pathways revealed a significant enrichment of functions related to host responses to the viral replication in cluster 2 and 3, with clathrin-mediated endocytosis (R-HSA-8856828) and vesicle-mediated transport (R-HSA-5653656) pathways specific of the former one (Figure 4D) and likely involved in vacuole formation and viral budding. Interestingly, similar to cluster 2 obtained at higher MOI, the expression profiles of the members of these clusters are similar to the dynamics of the viral protein levels across the kinetics. Clusters 4 (MOI 0.01) and 3 (MOI 0.001) were enriched in pathways and biological functions related to the host central metabolism with the metabolism of RNA and translation among the top enriched pathways. Despite the temporal expression profiles of the proteins belonging to these clusters are not characterised by major changes, their remodelling by the virus and a major effect on their abundance at earlier time points (<24 hours) cannot be excluded.

**Figure 3.**
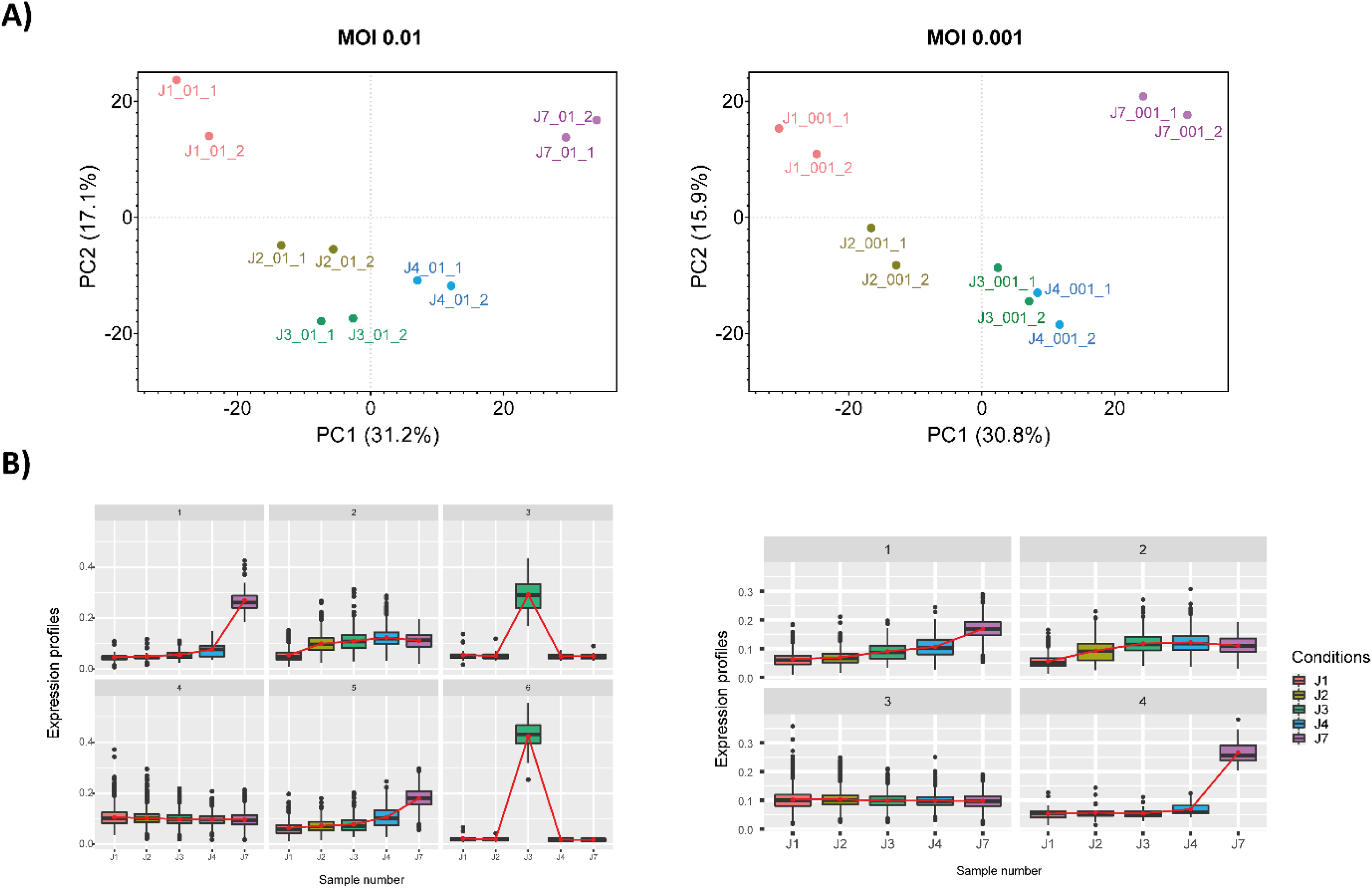
Host response upon SARS-CoV-2 infection. **(A)** Dimension reduction by principal component analysis (PCA) of the different time points at MOI 0.01 and 0.001, respectively. Normalised abundance of proteins with Spectral Count >3 was used as input. J1, J2, J3, J4 and J7 refer to the different time points analysed, Day 2, 3, 4 and 7, respectively. **(B)** Clusters of proteins showing similar expression profiles over time. J1, J2, J3, J4 and J7 refer to the different time points analysed, Day 2, 3, 4 and 7, respectively. The number on the top of each plot identifies the cluster.

**Figure 4.**
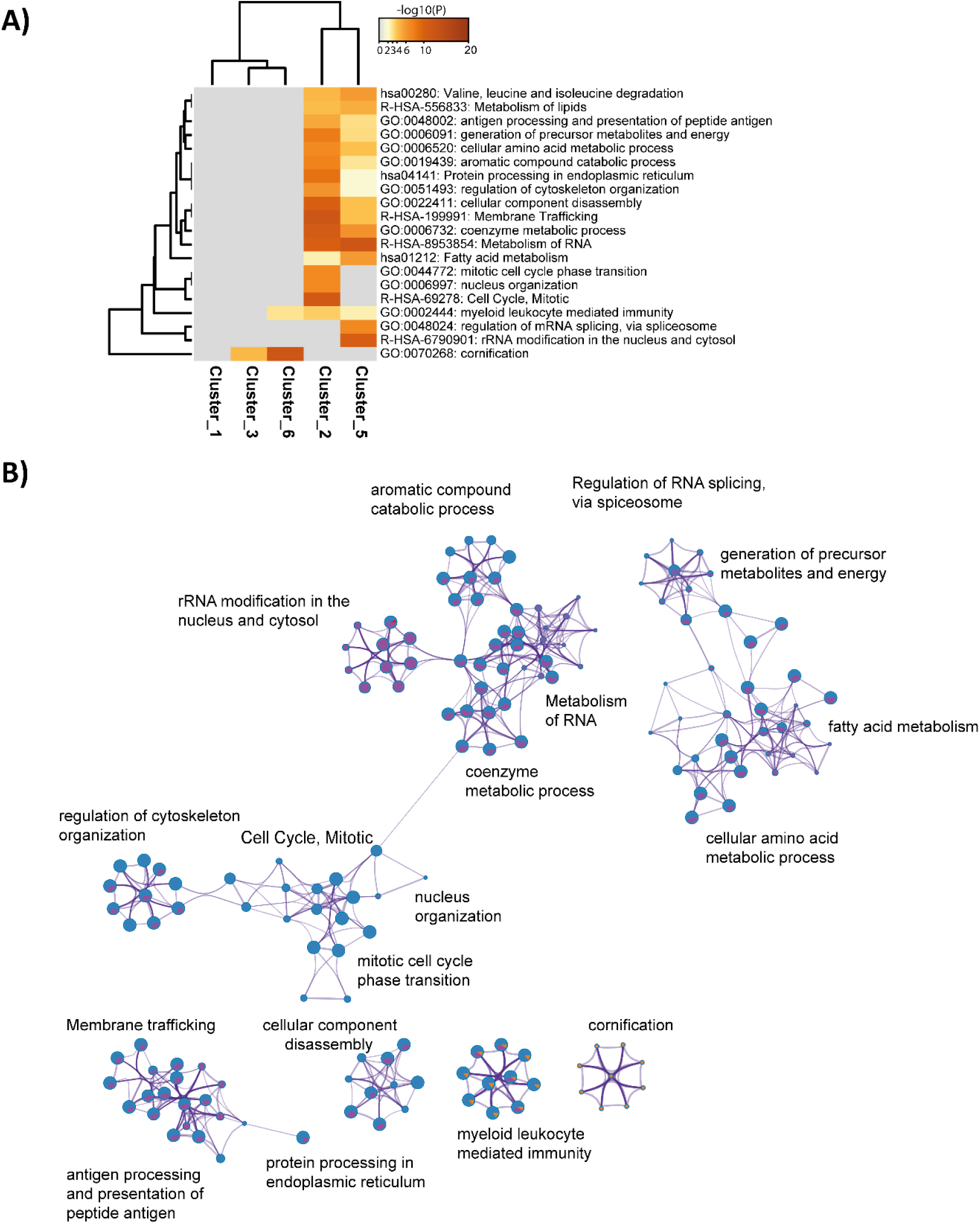
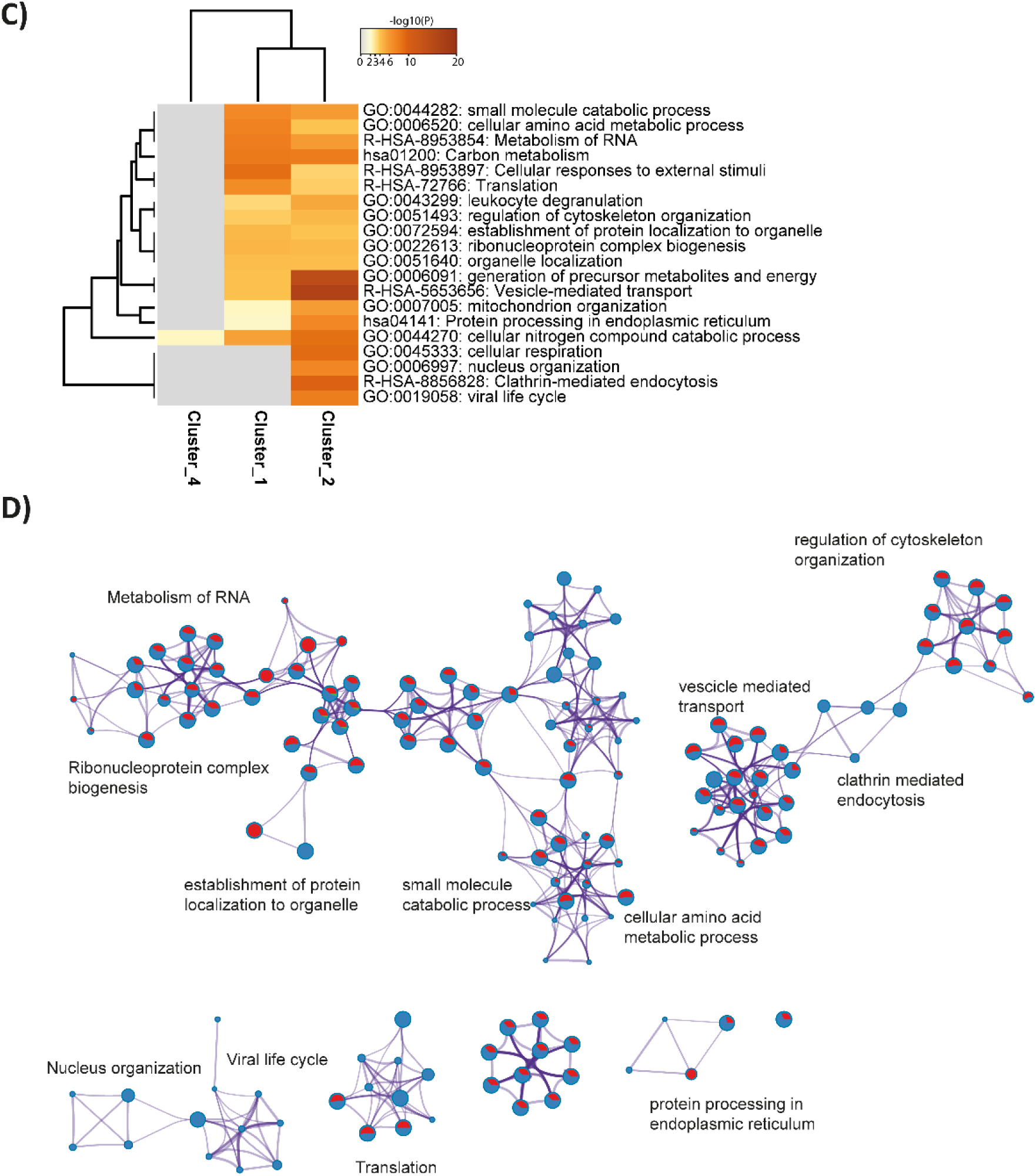
Host pathways dysregulated by the infection. **(A-C**) Heatmap showing the top enrichment clusters, one row per cluster, using a discrete color scale to represent statistical significance. Gray color indicates a lack of significance. MOI 0.01**(A)** and 0.001 **(C). (B-D)** Enrichment network visualization for results from the proteins present in each of the identified clusters. Nodes are represented by pie charts indicating their associations with each cluster. Color code represents the clusters. At MOI 0.01 **(B)**, red represents Cluster 1 while blue, green, purple and orange refer to Cluster 2, 3, 5 and 6, respectively. At MOI 0.001 **(D)**, red indicates Cluster 1 while blue and green refer to Cluster 2 and 4, respectively.

## Discussion

In the race to develop a vaccine for fighting the global spreading of SARS-CoV-2, different technology platforms have been evaluated [23]. In this context, the obtainment of vaccines based on inactivated or attenuated whole viral particles could be challenged in the finding of the right conditions for virus amplification. Virus yields from the dedicated cell culture systems could also represent a limitation. Concomitantly with vaccine development, inactivated virus particles are also of interest for testing real serology or screening neutralizing antibodies. Evidently, production of well-characterized active virus particles is also of interest for fundamental research purposes. Given the requirement for speed, here we evaluate the use of LC-MS/MS as a tool for guiding the optimisation of the conditions for SARS-CoV-2 whole viral particle antigen production.

The results presented here demonstrate the potential of our pipeline to profile virus production across time. In particular, by analysing the proteome of Vero cells infected with SARS-CoV-2 at two different MOI, it was possible to monitor changes in the levels of three SARS-CoV-2 structural proteins and three non-structural ones. Whilst as for other analyses [11,13] we could not detect peptides from protein E like. The lack of detection of other accessory proteins could be imputed to differences in samples processing with the protocol described here favouring simplified steps and speed while maintaining accuracy. Deeper analyses are envisaged for monitoring virus homogeneity during the different steps of viral production once the most permissive conditions will be established.

Remarkably, comparable profiles were obtained at the two tested MOI, with the profiles obtained at lower MOI slightly delayed and hence more insightful regarding the timing of the burst in the abundance of viral levels.

Overrepresented proteins were described by a larger variety of peptides. Interestingly, an increase of protein levels relied on the increasing abundance of the same set of peptides with only a few new sequences registered at late time points suggesting that absolute quantification of the virus could be obtained by targeted approaches by following early detectable peptides. Eventually tandem mass spectrometry proteotyping [24,25] could be also proposed to detect SARS-CoV-2 viruses.

Besides profiling virus production, our mass spectrometry analysis of the whole cell content provides insights regarding the cellular response to SARS-CoV–2 infection. Notwithstanding, while more detailed information is available regarding virus cell entry, increased understanding of the different steps in the SARS-CoV-2 replication cycle are needed [26]. To our knowledge, this is one of the first attempt to characterize cellular response to infection with SARS-CoV–2 in Vero E6 cells. Interestingly, such proteomics data can be acquired on different cell lines from humans and primates in order to define by comparative proteomics the common mechanisms of cell infection and the mechanisms specific of a given cell line.

The analysis of our proteomic data suggested substantial temporal remodelling of the host proteome over the time points. Functional enrichment analysis of clusters of proteins showing similar expression profiles highlight key pathways during virus replication providing a potential target for effective therapeutics against coronaviruses rises. Consistently with the pathways identified by Bojkova et al. [11] by using the human colon epithelial carcinoma cell line Caco–2 as system for the analysis of SARS-CoV-2 replication, we identified clusters of proteins increased by infection and enriched in RNA modifiers, such as spliceosome components, and carbon metabolism, further supporting the preliminary evidence that classifies splicing as an essential pathway for SARS-CoV–2 replication and potential therapeutic target. Additional clusters of proteins identified in our analysis highlighted the regulation of pathways critical for the virus life cycle like those involved in protein pre-processing in the endoplasmatic reticulum, vacuole formation and viral budding.

Here, we show that our pipeline based on LC-MS/MS analysis is a suitable tool for the characterisation of SARS-CoV-2 production. We, therefore, suggest that it could be of use in the optimisation of the condition for viral amplification to speed up the initial steps in favour of those that later on during the development process will require a more careful evaluation of effectiveness and safety.

Besides, peptide information described here provide sufficient information to enable a targeted analysis, opening the possibility of using mass spectrometry-based targeted approaches for the evaluation of critical aspects (*i.e.* quality and quantity) during the different steps of the virus purification processes. Furthermore, the characterized changes in cellular protein networks upon SARS-CoV-2 infection provided valuable insights that could be further explored and guide the identification of drug targets to address the pandemic caused by SARS-CoV-2.

We can anticipate that the same workflow could be successfully applied to expedite the characterisation of human organ-on-a-chip (Organ Chip) microfluidic culture devices used to obtain insights on the different steps of the virus life cycle as well as to study human disease pathogenesis [27] in response to infection by variants of SARS-Cov-2 under or not the addition of existing [28,29,30] and novel therapeutics.

## Author contributions

LG, FG, OP, LB, and JA conceptualized the stud design; FG, JCG, HB, SR, NB performed the experiments; LG, OP, DG, FG, LB and JA analysed the data; GM, KC, SD, MAR, GS, CF, AD, BAB, and CB contributed reagents and software development; LG and JA draft the manuscript. All authors read and approved the final manuscript.

## Acknowledgements

The authors are indebted to Dr Silvia Meschi (National Institute for Infectious Diseases “Lazzaro Spallanzani” IRCCS, via Portuense 292, 00149 Rome, Italia) for making the Human 2019-nCoV strain 2019-nCoV/Italy-INMI1 (008N-03893) available. The authors thank all their colleagues from Li2D (CEA) for strong support, as well as all those who make the experimental work possible while facing the Covid19 pandemic and lockdown.

## Disclosure statement

No potential conflict of interest was reported by the authors.

## Funding

This work was funded in part by the French Alternative Energies and Atomic Energy Commission (CEA), the French joint ministerial program of R&D against CBRNE threats, and the ANR program “Phylopeptidomics” (ANR-17-CE18-0023-01). This publication was supported by the European Virus Archive goes Global (EVAg) project that has received funding from the European Union’s Horizon 2020 research and innovation programme under grant agreement N°653316.

## Notes

https://www.ebi.ac.uk/pride/archive/projects/PXD018594

### Competing Interest Statement

The authors have declared no competing interest.

